# SARS-CoV-2 Envelope protein triggers depression and dysosmia via TLR2 mediated neuroinflammation

**DOI:** 10.1101/2023.01.15.524078

**Authors:** Wenliang Su, Jiahang Ju, Minghui Gu, Xinrui Wang, Shaozhuang Liu, Jiawen Yu, Dongliang Mu

## Abstract

**Background:** Depression and dysosmia have been regarded as the main neurological symptoms in COVID-19 patients, the mechanism of which remains unclear. Current studies have demonstrated that SARS-CoV-2 envelope protein served as a pro-inflammatory factor as sensed by Toll like receptor 2 (TLR2), suggesting the viral infection independent pathological feature of E protein. In this study, we aim to determine the role of E protein in depression, dysosmia and associated neuroinflammation in central nervous system (CNS).

**Methods:** Depression and olfactory function were observed in both female and male mice as receiving intracisternal injection of envelope protein. Immunohistochemistry was applied in conjunction with RT-PCR to assess the glial activation, blood-brain barrier status and mediators synthesis in cortex, hippocampus and olfactory bulb. TLR2 was pharmacologically blocked to determine its role in E protein related depression and dysosmia.

**Results:** Intracisternal injection of envelope protein evoked depression and dysosmia in both female and male mice. Immunohistochemistry suggested that envelope protein upregulated IBA1 and GFAP in cortex, hippocampus and olfactory bulb, while ZO-1 was downregulated. Moreover, IL-1β, TNF-α, IL-6, CCL2, MMP2 and CSF1 were upregulated in both cortex and hippocampus, whereas IL-1β, IL-6 and CCL2 were upregulated in olfactory bulb. Furtherly, inhibiting microglia, but not astrocyte, alleviated depression and dysosmia induced by envelope protein. Finally, RT-PCR and immunohistochemistry suggested that TLR2 was upregulated in cortex, hippocampus and olfactory bulb, the blocking of which mitigated depression and dysosmia induced by envelope protein.

**Conclusions:** Our study demonstrates that envelope protein could directly induce depression and dysosmia together with obvious neuroinflammation in CNS. TLR2 mediated depression and dysosmia induced by envelope protein, which could serve as a promising therapeutic target for neurological manifestation in COVID-19 patients.

## Introduction

The COVID-19 pandemic has dramatically impacted the world since the SARS-CoV-2 virus infection outbreak in 2019. Clinical symptoms of COVID-19 could involve several human body systems, apart from the most commonly known respiratory system symptoms. The COVID-19 infection can cause a series of neurological symptoms, including attention disorders, sleep disorders, short-term memory loss, seizures, strokes, headaches, dizziness, smell and taste loss, and neuropsychiatric symptoms such as anxiety and depression (Bartley et al., 2021; Chen et al., 2020; Mazza et al., 2020; Mazza et al., 2021; Saladino et al., 2020; Schou et al., 2021; Solomon et al., 2020; Su et al., 2020). Furthermore, COVID-19 patients’ brain MRI imaging showed the overall brain size, the prefrontal brain and parahippocampal gyrus gray matter thickness were decreased. In addition, the changes of tissue damage markers in the primary olfactory cortex related brain area were more significant compared with the negative control group (Smith, 2022). Further pathological observation revealed that brain samples from patients with COVID-19 showed lesions of SARS-CoV-2 infection and replication, and that the virus infected astrocytes (Crunfli et al., 2022). However, the relationship between the SARS-CoV-2 virus infecting and COVID-19 neurological symptoms is still unclear.

The SARS-CoV-2 virus has sixteen non-structural proteins (Nsp1-16) and four structural proteins, known as the membrane protein, spike protein, nucleocapsid protein, and envelope protein (E) (Brian and Baric, 2005). The E is a small integral protein built up by only 75 amino acids. However, E involves many processes in the virus’s life cycle, such as assembly, budding, envelope formation, and pathogenic mechanism (Schoeman and Fielding, 2019). The E highly expresses in infected cells during the virus’s replication cycle (Venkatagopalan et al., 2015). E plays a vital role in virus production and maturation. Evidence reveals that E-lacking recombinant coronavirus showed significantly decreased virus titer and offspring with impaired virus maturation or insufficient production and proliferation (DeDiego et al., 2007; Ortego et al., 2007). Meanwhile, E, as a multifunctional protein, not only acts as a structural protein in the virus capsid and participates in virus assembly but also acts as a virulent toxin and participates in the pathogenesis of the virus (Cao et al., 2021; Mandala et al., 2020).

Previous evidence reported that the E protein of SARS-CoV-2 could bind to the Toll-like receptors (TLRs) and drive pulmonary inflammation (Zheng et al., 2021). At the same time, the E protein is also necessary for inflammatory cytokines release during coronavirus infection. Using E protein stimulated human peripheral blood mononuclear cells, it was found that the E protein could interact with human TLR2 receptors and induce the expression of tumor necrosis factor, interferon-γ, interleukin 6, and interleukin 1β. SARS-CoV-1 strain lacking E protein could not activate the NF- κB signal transduction pathway and significantly decreased inflammatory cytokines production (DeDiego Marta et al., 2014). Combined with the neurological SARS-CoV-2 infection and the vital role of E protein-TLRs interaction in the virus pathogenesis, we hypothesize that the COVID-19 patients’ dysosmia and depression symptoms might be mediated by the interaction of E protein with TLRs of glial cells.

## Methods

### Animals

Wild type C57BL/6 male and female mice (20–25 g; 6–8 weeks), purchased from the National Institutes for Food and Drug Control in China, were used in this study. Animals were housed under a 12-h light/12-h dark cycle with ad libitum access to food and water.

### Sucrose preference test (SPT)

The mice were put into the cages with two-bottle drinking setting for 48h (one bottle for water, and the other for 1.5% sucrose solution). In addition, the position of the two bottles was switched at 24h. The water access was then deprived for 24h to the tested mice. The two-bottle drinking setting was provided once again to the tested mice for 2h, during which the position of the two bottles was switched at 1h. The ratio of sucrose solution consumed to the total fluid intake was determined as the sucrose preference (Liu et al., 2021).

### Tail suspension test (TST)

The TST was performed as previously described. Briefly, each mouse was suspended by its tail with a short adhesive tape connected to a load cell that transmitted a signal corresponding to activity. The total test time was 6□min. After setting a low threshold, the duration of immobility was recorded and analyzed by tail suspension software (SOF-821, Med Associates) (Liu et al., 2021).

### Forced swimming test (FST)

Mice were individually placed in a beaker (height: 19□cm; diameter: 14□cm) containing 14□cm of water (23□±□2□°C). The total test duration was 6□min. The process was videotaped, and the immobility time in the last 4□min was scored by an experienced observer blinded to the experimental treatment. Floating or only slight movement to maintain balance was considered as the immobility (Liu et al., 2021).

### Olfactory measurement

At the first day, the mice were trained for 3 min in a cage which contained sunflower seeds in four corners. Then, the mice were trained for 3 min again, but sunflower seeds were placed only in one corner. After training, the mice were separated from their dam and food was withheld for one day. Three 3-min trials were conducted at day 3. In trials 1 and 2, one sunflower seed was placed in the cage on bedding, each time in a different corner. In trial 3, a seed was buried at the center of the cage under the bedding. Latency to find the food was recorded. If mice were unable to find a seed within 3 min, the latency to find food was recorded as 180 s. Shorter latency to find buried sunflower seed indicates better olfaction (Hung et al., 2018).

### Intracisternal injection

To observe the effects of E protein in CNS, mice received intracisternal injection of E protein. Briefly, E protein (0.2 μg/μl buffered in PBS, 5 μl) (ENN-C5128, Acro Biosystems) or Vehicle (PBS, 5 μl) was injected slowly into the cisterna magna with a specially-made length-limited syringe (Ju et al., 2022). To inhibiting microglia and astrocyte, minocycline (a microglial inhibitor, 5 μg in 5 μl PBS, daily delivered for constructive 5 days prior to E protein application) (M9511, sigma) and L-α-aminoadipate (LAA, an astroglial toxin, 50 nmol buffered in 1N HCl and further diluted in PBS, delivered 1 hour prior to E protein application) (A7275, sigma) were applied via intracisternal injection (Chen et al., 2018; Zhao et al., 2021). To specifically block TLR2, C29 (a specific TLR2 inhibitor, 50 mg/kg, buffered in PBS containing 10% DMSO, delivered 1 hour prior to E protein application) inhibitor was also injected into cistern by syringe. The timeline of drug delivery, behavior test and tissue collection had been showed in Figure 1.

**Figure 1.**
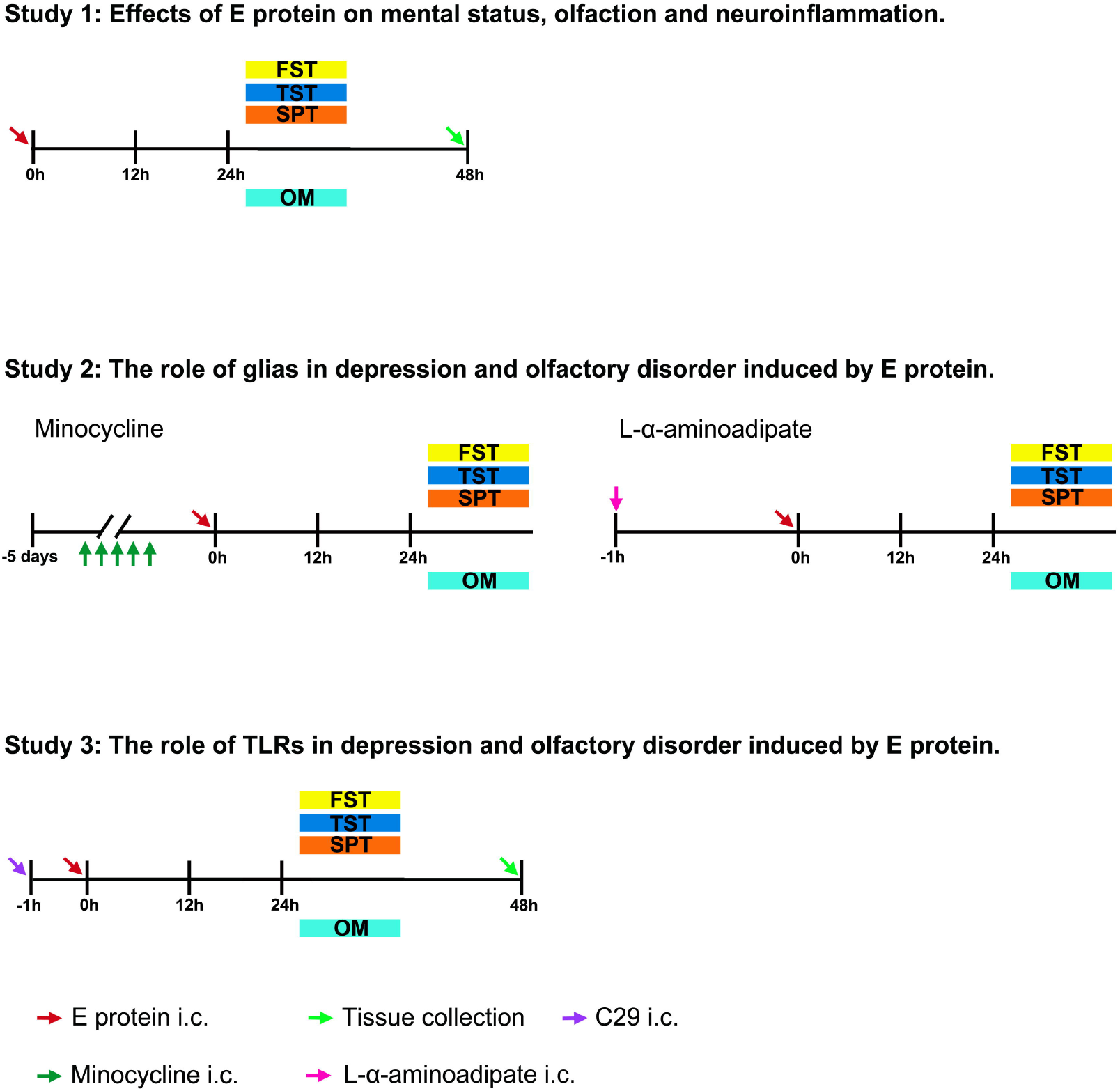
The timeline of E protein injection, drug delivery and behavioral test.

### Immunofluorescence

Immunofluorescence was performed as previously described. Briefly, mice were anesthetized with intraperitoneal sodium pentobarbital injection (40 mg/kg) under aseptic condition and then transcardially perfused with ice-cold PBS followed by ice-cold paraformaldehyde. The whole brain from each tested mouse was collected, post-fixed in 4% paraformaldehyde at 4°C for 2h, and then dehydrated in 30% sucrose at 4 °C. Then, these brain tissues were embedded in OCT (Tissue-Tek, Japan) and serially sectioned in a cryostat (Leica 2000, Germany) into 30-μm-thickness slices. The tissue sections were blocked with 5% donkey serum containing 0.3% Triton X-100 for 1h, and then primary antibodies were incubated at 4°C overnight in a wet box. After washed in PBS buffer, secondary antibodies were incubated for 1h at room temperature (Primary antibodies list: ZO-1, ab 221547, abcam, 1:200; IBA1, ab5076, abcam, 1:1000; GFAP, ab53554, abcam, 1:1000; TLR2, ab209216, abcam, 1:200. Secondary antibody list: Donkey anti-Rabbit 488, A32790, Invitrogen, 1:500; Donkey anti-Mouse 488, A-21202, Invitrogen, 1:500; Donkey anti-Goat 488, ab150129, abcam, 1:500). The slides were then coverslipped by Mounting Medium solution containing DAPI (ZSJB-Bio, Beijing, China). Images were captured by laser confocal microscopic imaging system under the same settings (TCS-SP8 STED 3X, Leica, Germany). The quantification for immunofluorescence staining referred to previous studies (Su et al., 2021). Briefly, at least 10 sections from 3 randomly selected mice on each group were examined. The positive area of IBA1, GFAP, ZO-1 and TLR2 staining was measured with Image J.

### Quantitative RT-PCR

Total RNAs of cortex, hippocampus and olfactory bulb were extracted by Trizol reagent (CW-bio, Beijing, China), which were then reversely transcribed by RT Master Mix according to the instruction (Takara, Japan). Finally, qRT-PCR was performed using a CFX96™ Real-Time PCR Detection System (Bio-Rad, California, USA) with TB green premix Ex Taq (Takara, Japan). The primers used were listed in Table S1.

### Statistical analysis

Data values were presented as means with standard errors (mean□± □SEM). Statistical analyses were performed by Graph Pad software. Shapiro-Wilk test was applied to determine the normality for parametric test. The student’s t test was used to determine statistical difference between two groups. The criterion for statistical significance was a value of *p<0*.*05*.

## Results

### E protein induced depression and dysosmia in female and male mice

To identify the neuropathological effects of E protein, we applied i.c. injection of E protein and evaluated the mental and olfactory status. The SPT, TST, and FST tests indicated that E protein in the central nervous system significantly induced depression in both female and male mice (Figure 2A-C). By the way, olfactory measurement showed that E protein increased the costing time for finding the target food in both female and male mice, suggesting that E protein evoked smelling failure (Figure 2D). Collectively, we observed the typical depression and olfactory disturbance symptoms in COVID-19 patients as delivering E protein to central nervous system.

**Figure 2.**
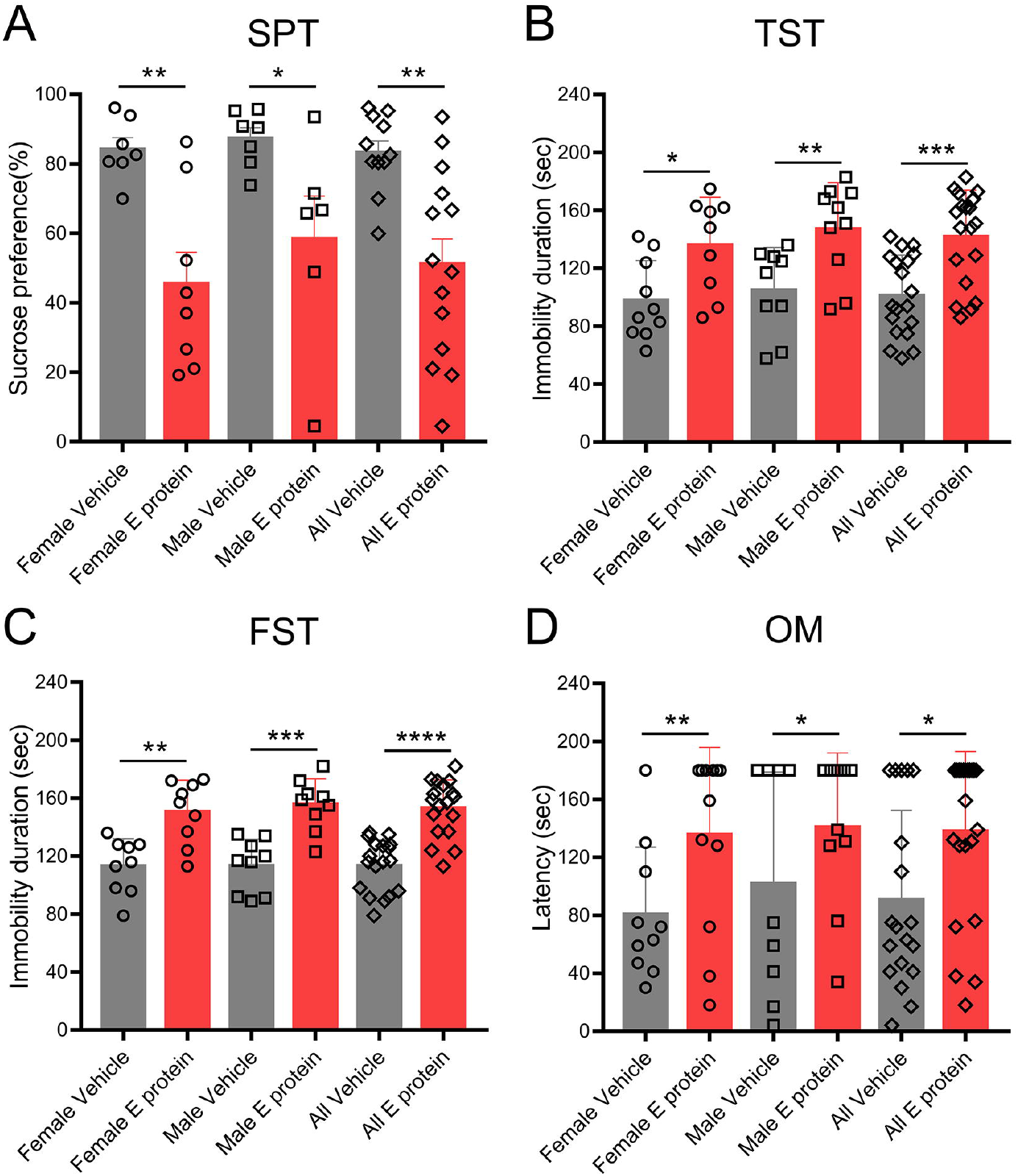
The SPT, TST, FST, and olfactory measurement for mice receiving intracisternal injection of E protein. **A**. The percentage of sucrose water consumption for female and male mice receiving Vehicle or E protein injection. **B**. The immobility duration in TST test for female and male mice receiving Vehicle or E protein intracisternal injection. **C**. The immobility duration in FST test for female and male mice receiving Vehicle or E protein intracisternal injection. **D**. The latency time for discovering the sunflower seed in olfactory measurement. \**p<0*.*05*, \*\**p<0*.*01*, \*\*\**p<0*.*001*, \*\*\*\**p<0*.*0001*; student’s t test.

### E protein triggered neuroinflammation and blood-brain barrier damage in cortex, hippocampus and olfactory bulb

Neuroinflammation has been regarded as a key cause of depressive and olfactory disorder, therefore we further systemically assessed the neuroinflammatory status in relative cortex, hippocampus and olfactory bulb regions. The IF staining for microglia marker IBA1 suggested significant microglia proliferation induced by E protein in cortex (Figure 3A-B), hippocampus (CA1, CA3 and DG regions) (Figure 3C-F), and olfactory bulb (Figure 3G-H) in both male and female mice. Meanwhile, GFAP, the astrocytes marker, was also upregulated by E protein in cortex (Figure 4A-B), hippocampus (CA1, CA3 and DG regions) (Figure 4C-F), and olfactory bulb (Figure 4G-H). These above results indicated that E protein triggered significant glial activation in cortex, hippocampus and olfactory bulb regions. The blood-brain barrier (BBB) destruction plays a critical role in neuroinflammation formation; therefore, we also tested the expression of tight junction marker ZO-1 to evaluate BBB status. The IF staining showed significant decrease of ZO-1 expression in cortex (Figure 5A-B), hippocampus (CA1, CA3 and DG regions) (Figure 5C-F), and olfactory bulb (Figure 5G-H) induced by E protein injection in both female and male mice. Furtherly, we screened the typical neuroinflammatory mediators in relative brain areas. The RT-PCR results showed that IL-1β, TNF-α, IL-6, CCL2, MMP2 and CSF1 mRNA expression were upregulated in cortex and hippocampus. Meanwhile, the IL-1β, IL-6 and CCL2 mRNA expression were upregulated in olfactory blob (Figure 6). Together, the above results suggested the occurrence of neuroinflammation in depression and dysosmia relative brain regions induced by E protein.

**Figure 3.**
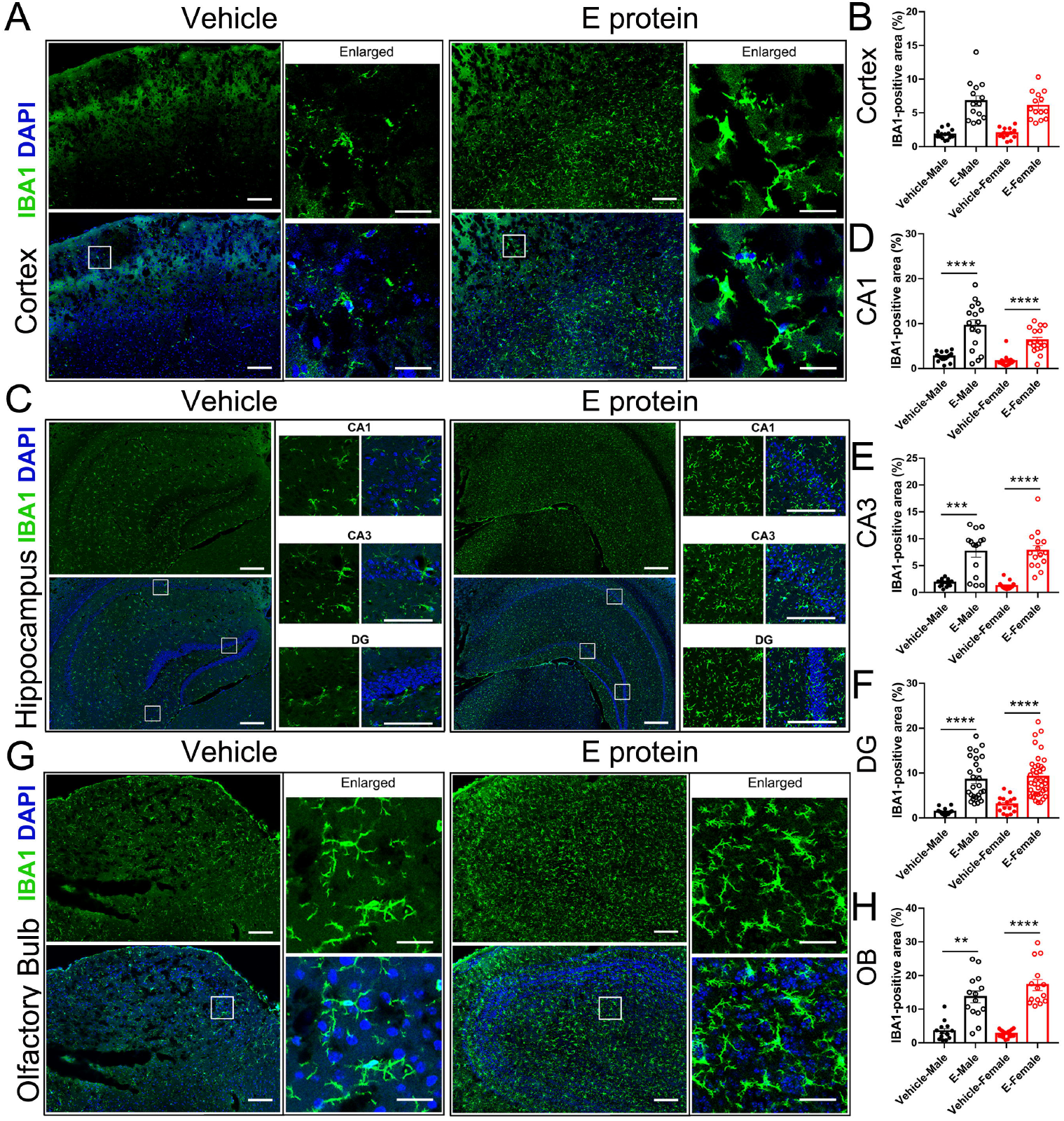
Intracisternal injection of E protein upregulated IBA1 in cortex, hippocampus and olfactory bulb. **A**. Representative images of IBA1 expression in cortex from mice receiving Vehicle or E protein injection. **B**. Fluorescence area analysis showed that E protein significantly upregulated IBA1 expression in cortex. **C**. Representative images of IBA1 expression in hippocampus (CA1, CA3 and DG regions) from mice receiving Vehicle or E protein injection. **D-F**. Fluorescence area analysis showed that E protein significantly upregulated IBA1 expression in CA1, CA3 and DG regions of hippocampus. **G**. Representative images of IBA1 expression in olfactory bulb from mice receiving Vehicle or E protein injection. **H**. Fluorescence area analysis showed that E protein significantly upregulated IBA1 expression in olfactory bulb. \**p<0*.*05*, \*\**p<0*.*01*, \*\*\**p<0*.*001*, \*\*\*\**p<0*.*0001*; student’s t test.

**Figure 4.**
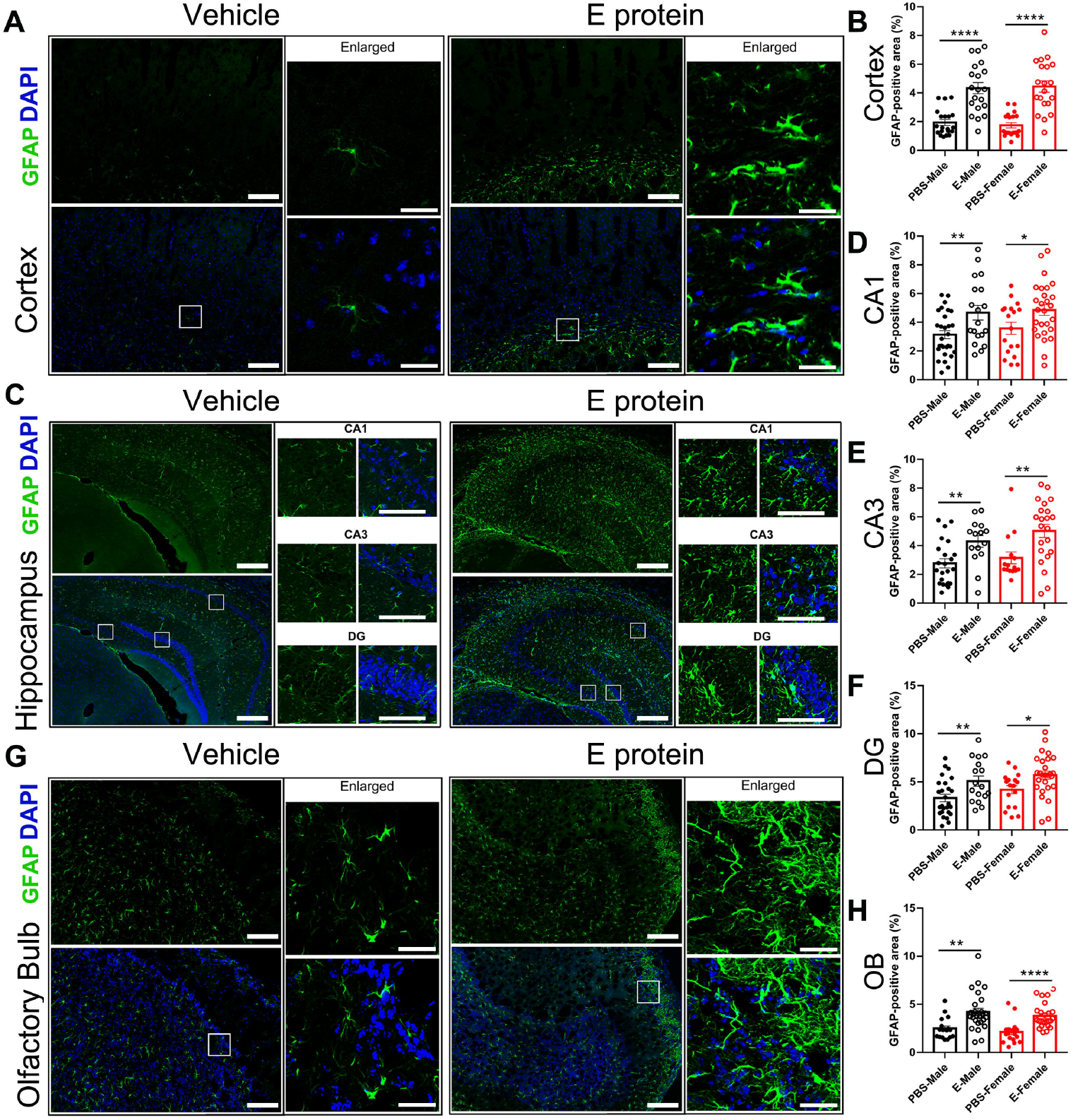
Intracisternal injection of E protein upregulated GFAP in cortex, hippocampus and olfactory bulb. **A**. Representative images of GFAP expression in cortex from mice receiving Vehicle or E protein injection. **B**. Fluorescence area analysis showed that E protein significantly upregulated GFAP expression in cortex. **C**. Representative images of GFAP expression in hippocampus (CA1, CA3 and DG regions) from mice receiving Vehicle or E protein injection. **D-F**. Fluorescence area analysis showed that E protein significantly upregulated GFAP expression in CA1, CA3 and DG regions of hippocampus. **G**. Representative images of GFAP expression in olfactory bulb from mice receiving Vehicle or E protein injection. **H**. Fluorescence area analysis showed that E protein significantly upregulated GFAP expression in olfactory bulb. \**p<0*.*05*, \*\**p<0*.*01*, \*\*\**p<0*.*001*, \*\*\*\**p<0*.*0001*; student’s t test.

**Figure 5.**
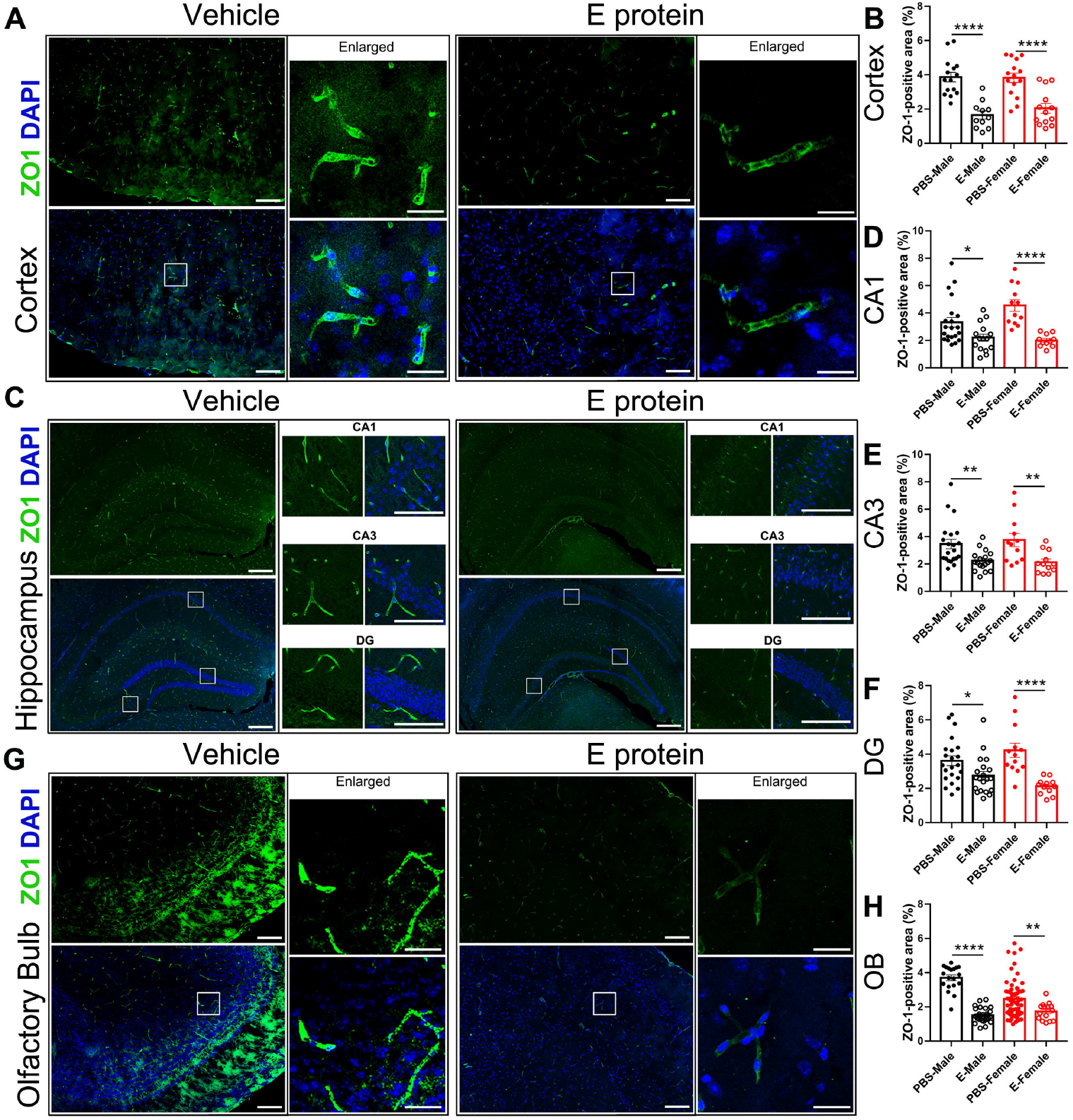
Intracisternal injection of E protein downregulated ZO-1 in cortex, hippocampus and olfactory bulb. **A**. Representative images of ZO-1 expression in cortex from mice receiving Vehicle or E protein injection. **B**. Fluorescence area analysis showed that E protein significantly downregulated ZO-1 expression in cortex. **C**. Representative images of ZO-1 expression in hippocampus (CA1, CA3 and DG regions) from mice receiving Vehicle or E protein injection. **D-F**. Fluorescence area analysis showed that E protein significantly downregulated ZO-1 expression in CA1, CA3 and DG regions of hippocampus. **G**. Representative images of ZO-1 expression in olfactory bulb from mice receiving Vehicle or E protein injection. **H**. Fluorescence area analysis showed that E protein significantly downregulated ZO-1 expression in olfactory bulb. \**p<0*.*05*, \*\**p<0*.*01*, \*\*\**p<0*.*001*, \*\*\*\**p<0*.*0001*; student’s t test.

**Figure 6.**
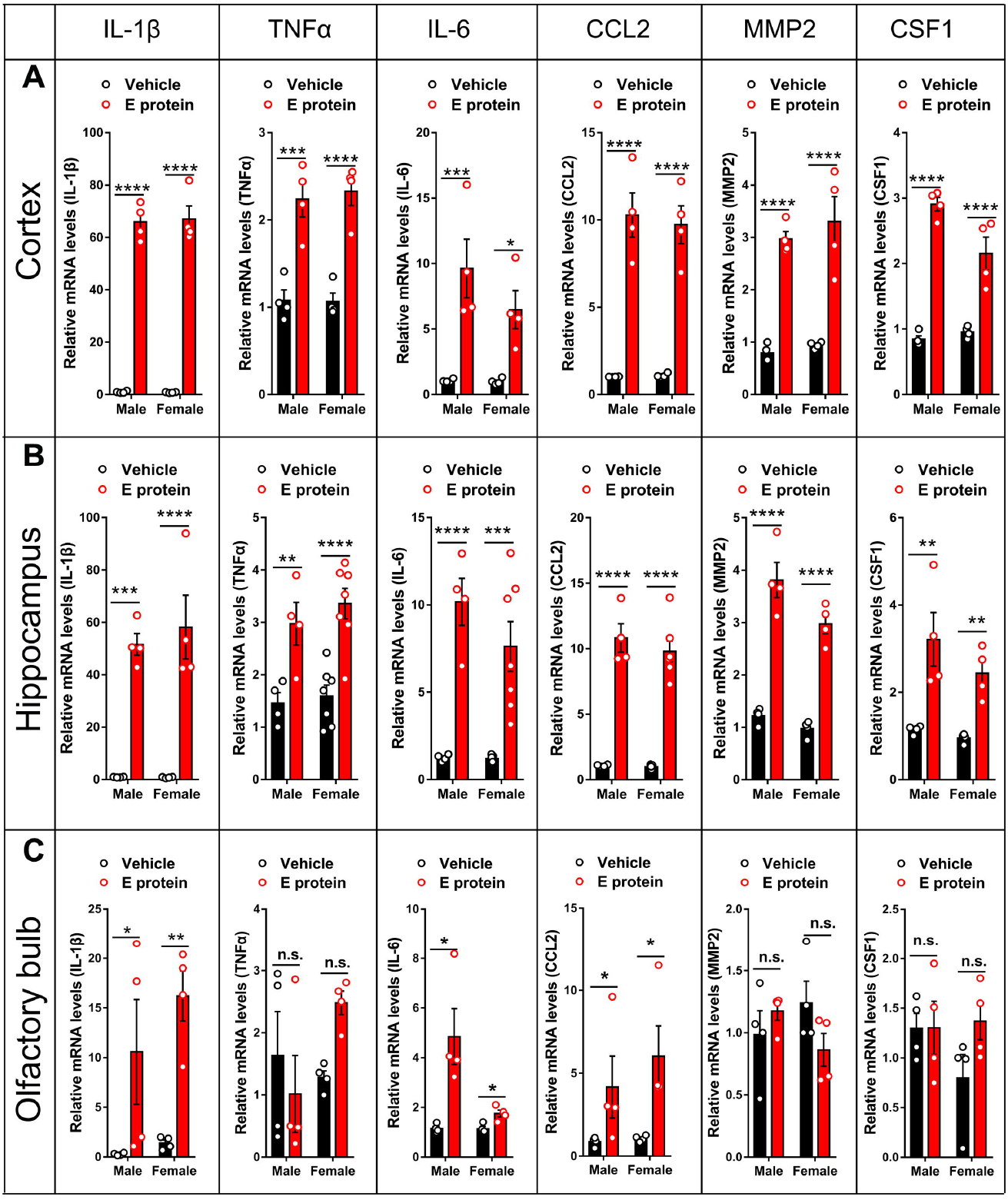
Intracisternal injection of E protein induced neuroinflammatory mediators in cortex, hippocampus and olfactory bulb. **A**. qRT-PCR analysis showed that intracisternal injection of E protein upregulated IL-1β, TNF-α, IL-6, CCL2, MMP2 and CSF1 in cortex. **B**. qRT-PCR analysis showed that intracisternal injection of E protein upregulated IL-1β, TNF-α, IL-6, CCL2, MMP2 and CSF1 in hippocampus. **C**. qRT-PCR analysis showed that intracisternal injection of E protein upregulated IL-1β, IL-6, and CCL2 in olfactory bulb. \**p<0*.*05*, \*\**p<0*.*01*, \*\*\**p<0*.*001*, \*\*\*\**p<0*.*0001*; student’s t test.

### Inhibiting microglia mitigated depression and dysosmia induced by E protein

Both microglia and astrocyte are vital elements in neuroinflammation and neuropathological process. To further investigate the contribution of microglia and astrocyte in E-induced depression and dysosmia symptoms, we used minocycline and L-α-Aminoadipate to inhibit microglia and astrocyte respectively before E protein administration. The SPT, TST, and FST tests showed that microglia inhibition by minocycline could successfully alleviate E-induced depression and olfactory disorder in female and male mice (Figure 7A-D). However, the astrocyte inhibition by L-α-Aminoadipate failed to recover the depression and olfactory disorder in mice, at least in at our observed time point (Figure 7E-H). Taken together, those results indicated that it was microglia rather than astrocyte that mediated E-induced depression and dysosmia.

**Figure 7.**
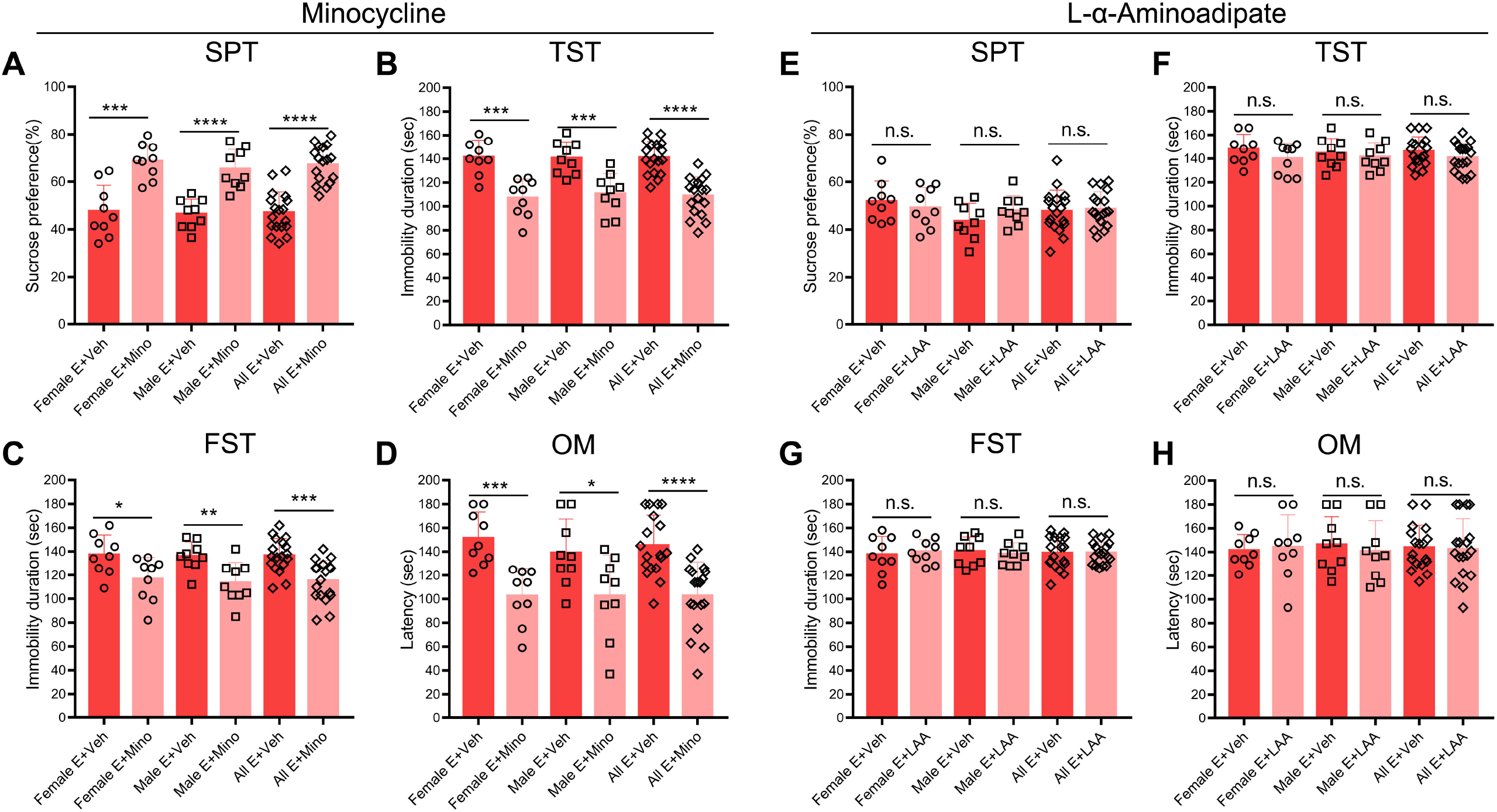
Inhibiting microglia alleviated depression and dysosmia induced by E protein. **A**. The percentage of sucrose water consumption for female and male mice receiving E protein injection as treated by Minocycline or Vehicle. **B**. The immobility duration in TST test for female and male mice receiving E protein injection as treated by Minocycline or Vehicle. **C**. The immobility duration in FST test for female and male mice receiving E protein injection as treated by Minocycline or Vehicle. **D**. The latency time for discovering the sunflower seed for female and male mice receiving E protein injection as treated by Minocycline or Vehicle. \**p<0*.*05*, \*\**p<0*.*01*, \*\*\**p<0*.*001*, \*\*\*\**p<0*.*0001*; student’s t test for A-D. **E**. The percentage of sucrose water consumption for female and male mice receiving E protein injection as treated by L-α-aminoadipate or Vehicle. **F**. The immobility duration in TST test for female and male mice receiving E protein injection as treated by L-α-aminoadipate or Vehicle. **G**. The immobility duration in FST test for female and male mice receiving E protein injection as treated by L-α-aminoadipate or Vehicle. **H**. The latency time for discovering the sunflower seed for female and male mice receiving E protein injection as treated by L-α-aminoadipate or Vehicle. n.s., no significance, student’s t test for E-H.

### E protein upregulated TLRs in cortex, hippocampus and olfactory bulb

As an exogenous protein, E protein might be recognized by Toll-like receptors (TLRs), a group of classic pattern recognition receptors. To identify the possible interaction between TLRs and E protein, we investigated the TLRs mRNA expression in associated brain regions. The RT-PCR results showed that TLR2, TLR3, TLR5, and TLR8 mRNA expression were upregulated in cortex (Figure 8A). Meanwhile, the E protein injection also significantly upregulated TLR2, TLR3, TLR4, TLR5 and TLR8 in the hippocampus (Figure 8B). In the olfactory bulb, the TLR2, TLR7 and TLR8 mRNA expression level were upregulated by E protein injection (Figure 8C). Those above results indicated that E protein might take effect via activating the TLRs dependent signal pathway. Given that TLR2 was upregulated among three brain regions and its crucial role in microglia function, we further investigated the distribution and contribution to E-related depression and dysosmia of TLR2.

**Figure 8.**
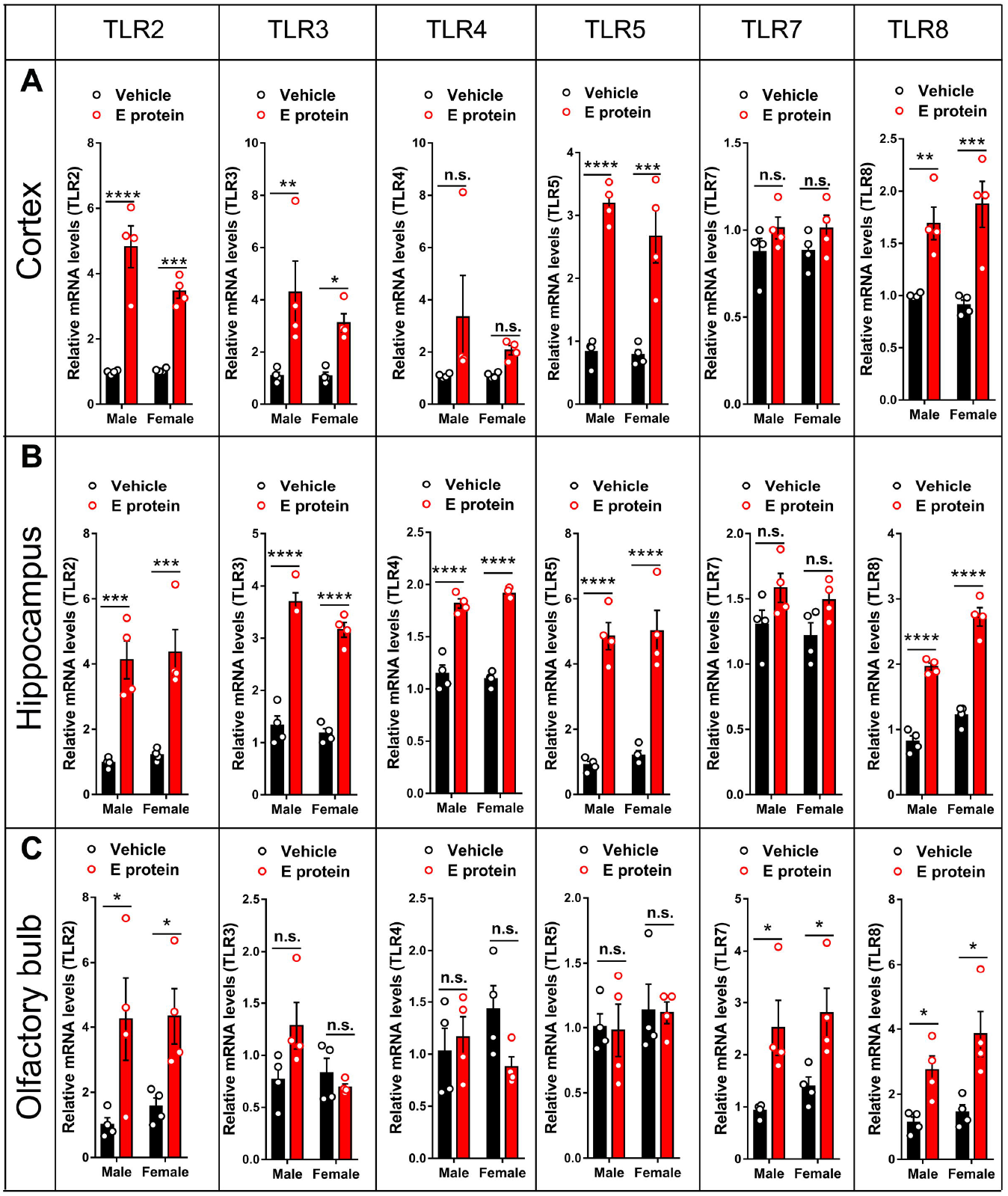
Intracisternal injection of E protein upregulated TLRs in cortex, hippocampus and olfactory bulb. **A**. qRT-PCR analysis showed that intracisternal injection of E protein upregulated TLR2, TLR3, TLR5 and TLR8 in cortex. **B**. qRT-PCR analysis showed that intracisternal injection of E protein upregulated TLR2, TLR3, TLR4, TLR5, and TLR8 in hippocampus. **C**. qRT-PCR analysis showed that intracisternal injection of E protein upregulated TLR2, TLR7, and TLR8 in olfactory bulb. \**p<0*.*05*, \*\**p<0*.*01*, \*\*\**p<0*.*001*, \*\*\*\**p<0*.*0001*; student’s t test.

### Blocking TLR2 alleviated depression and dysosmia induced by E protein

We supposed that E protein triggered neuroinflammation and evoked depression and dysosmia symptoms via TLR2 in CNS. Firstly, the IF staining of TLR2 suggested that E protein application to CNS significantly upregulated TLR2 expression in cortex (Figure 9A-B), hippocampus (CA1, CA3 and DG regions) (Figure 9C-F), and olfactory bulb (Figure 9G-H). Furthermore, we pharmacologically blocked TLR2 and measured the behavioral outcomes. The SPT, TST, and FST tests indicated the depression induced by E protein was alleviated by C29, a specific TLR2 blocker (Figure 10A-C). The shortened latency time in OM test suggested the olfactory disorder in mice was also relieved by C29 (Figure 10D). Above all, those results revealed that blocking TLR2 could alleviate E-induced dysosimia and depression.

**Figure 9.**
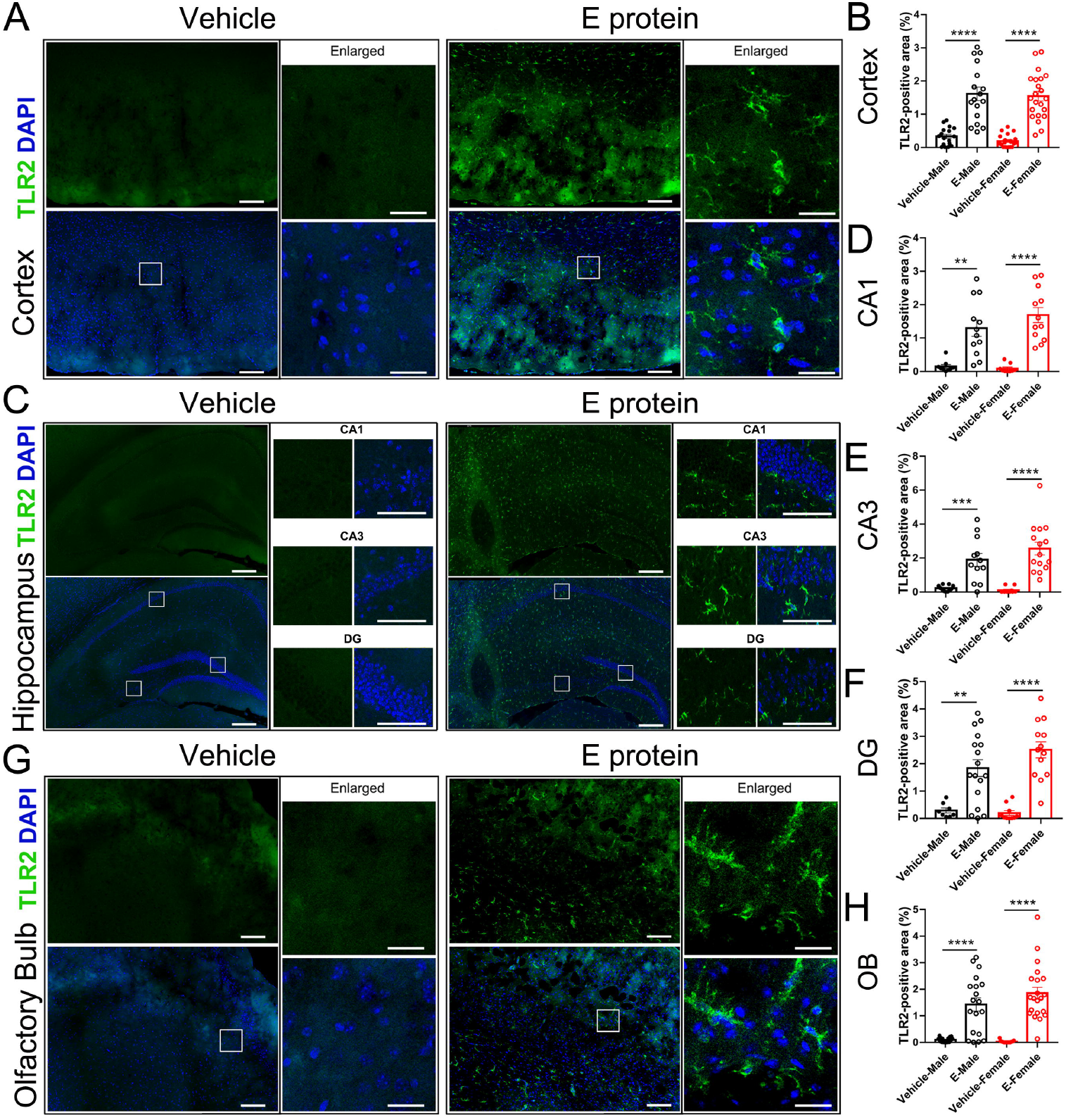
Intracisternal injection of E protein upregulated TLR2 in cortex, hippocampus and olfactory bulb. **A**. Representative images of TLR2 expression in cortex from mice receiving Vehicle or E protein injection. **B**. Fluorescence area analysis showed that E protein significantly upregulated TLR2 expression in cortex. **C**. Representative images of TLR2 expression in hippocampus (CA1, CA3 and DG regions) from mice receiving Vehicle or E protein injection. **D-F**. Fluorescence area analysis showed that E protein significantly upregulated TLR2 expression in CA1, CA3 and DG regions of hippocampus. **G**. Representative images of TLR2 expression in olfactory bulb from mice receiving Vehicle or E protein injection. **H**. Fluorescence area analysis showed that E protein significantly upregulated TLR2 expression in olfactory bulb.

**Figure 10.**
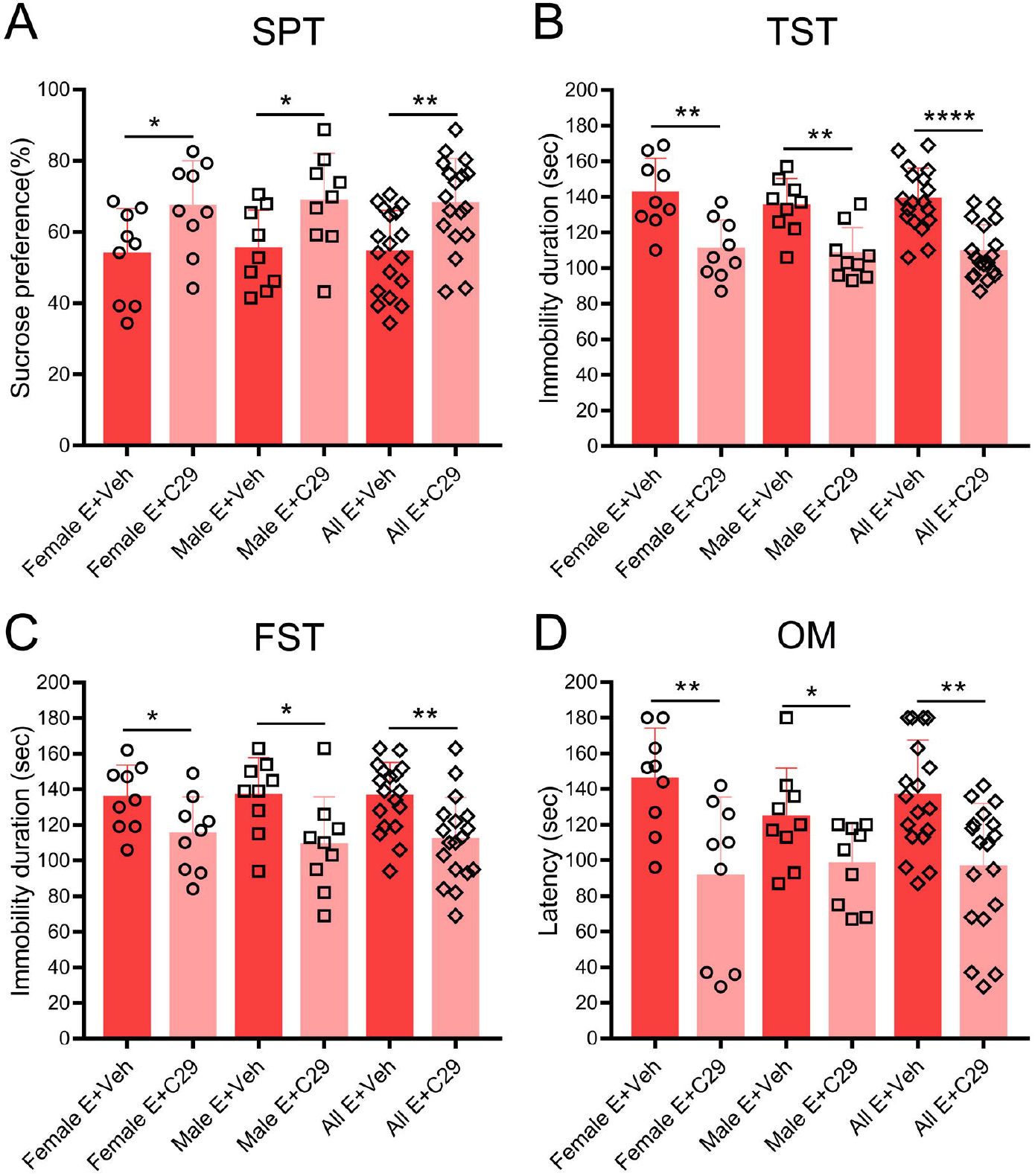
Blocking TLR2 alleviated depression and dysosmia induced by E protein. **A**. The percentage of sucrose water consumption for female and male mice receiving E protein injection as treated by C29 or Veh. **B**. The immobility duration in TST test for female and male mice receiving E protein injection as treated by C29 or Veh. **C**. The immobility duration in FST test for female and male mice receiving E protein injection as treated by C29 or Veh. **D**. The latency time for discovering the sunflower seed for female and male mice receiving E protein injection as treated by C29 or Veh. \**p<0*.*05*, \*\**p<0*.*01*, \*\*\**p<0*.*001*, \*\*\*\**p<0*.*0001*; student’s t test.

## Discussion

Patients with COVID-19 suffer from various neurological and psychiatric symptoms such as dysosmia and depression (Baig, 2020; Pezzini and Padovani, 2020). The cellular and molecular mechanisms are still elusive, though. The recent discovery provides evidence in support of the notion that the SARS-CoV-2 envelope (E) protein may cause a neuroinflammatory response separate from viral infection (Ju et al., 2022; Zheng et al., 2021). Given these proinflammatory properties of the E protein and that neuroinflammation often leads to olfactory dysfunction and mood disorder, we examined the ability of the E protein to trigger inflammatory responses in CNS in vivo and the behavioral sequelae of that response. We demonstrated that intracisternal injection of E protein led to depression and dysosmia as well as neuroinflammation in CNS. Our results suggest that several neurological and psychiatric disorders observed in COVID-19 patients can be mediated by the E protein during SARS-CoV-2 infection independent of viral entry.

Hyperactivation of the neuroimmune system is closely related to mood disorders and olfactory dysfunction. There is mounting evidence that microglia play an etiological role in this process (Frick et al., 2013; Kim et al., 2018). Depression is considered a microglia-associated disorder. Suicidal people and depressive patients exhibit notably elevated microglia activation (Schnieder et al., 2014; Steiner et al., 2008). Several depression-related brain regions have shown sustained microglial activation exhibiting high amounts of pro-inflammatory cytokines (Liu et al., 2019; Sugama et al., 2009; Zhang et al., 2016b). Microglial cells are highly concentrated in hippocampus, particularly in the CA1 region, and activation of these cells has been linked to the pathophysiology of major depressive disorder (Walker et al., 2013). Depressive-like behaviors were reduced by inhibiting microglial activation and neuroinflammation (Zhao et al., 2019). Some clinical antidepressants, such as nonsteroidal anti-inflammatory drugs, appear to reduce the symptoms of depression by preventing the activation of microglia (Jiang et al., 2019; Köhler et al., 2014; Zhong et al., 2019). In addition, CNS inflammation is one of the etiologies of olfactory disorders (Seo et al., 2018). Olfaction is a crucial sense controlled by the olfactory epithelium’s perception of odor molecules, which is then transported to the olfactory bulb (OB) via olfactory nerves and processed in the brain (Mori et al., 1999). There are lots of microglia in the OB. Microgliosis is frequently observed in the OB of olfactory dysfunction animals (Hovakimyan et al., 2013; Seo et al., 2016). Olfactory impairments may be caused by the microglial reaction in the OB which leads to the loss of neuroblasts (Seo et al., 2014).

The E protein induces potent inflammatory responses as a virulence factor (Xia et al., 2021). Microglia and astrocytes were activated by intracisternal injection of the E protein. SARS-CoV-2 virus caused microglial activation and astrogliosis, according to a postmortem case study (Matschke et al., 2020). Microglia and astrocytes are CNS resident cells and play a crucial role in homeostasis and neuroinflammation (Liu et al., 2020). As the innate immune cells in the brain, microglia are more sensitive to pathogens than astrocytes and serve as the main mediators of neuroinflammation (Garland et al., 2022). In response to CNS injury, microglial activation is essential for host defense and neuron survival (Polazzi and Contestabile, 2002). However, persistent activation and dysregulation of microglia may result in deleterious and neurotoxic consequences by overproduction of a variety of cytotoxic factors such as TNF-α (Sawada et al., 1989). Toxins and pathological stimuli injure neurons, which is enhanced and amplified by overactive microglia (Block and Hong, 2005). This then further causes more extensive damage to nearby neurons and ultimately promotes pathogenic outcomes (Teismann et al., 2003; Vasek et al., 2016).

On the other hand, both microglia and astrocytes can release proinflammatory mediators upon activation. In line with these findings, the E protein showed elevated expression of pro-inflammatory cytokines IL-1β and IL-6 in cortex, hippocampus, and OB. Recent findings showed that increased levels of IL-1β and IL-6 were discovered in the cerebral fluid of patients with COVID-19 infection and neurological symptoms (Normandin et al., 2021; Pilotto et al., 2021). Importantly, IL-1β and IL-6 have been identified as targets for alleviating the clinical condition, because they are thought to produce detrimental effects including the neurotoxic effect and inflammation. IL-1β has various biological activities. It increases neuronal apoptosis as well as neuronal loss by NMDA-evoked, glia-triggered, and/or proNGF-mediated pathways (Choi and Friedman, 2014; Long-Smith et al., 2010; Lu et al., 2005; Ye et al., 2013). IL-6 also has been linked to neuronal cell death (Conroy et al., 2004). Furthermore, we observed that the E protein reduced the expression of ZO-1 in various brain regions. IL-1β and IL-6 have been shown to enhance blood-brain barrier (BBB) permeability through cytokines-induced tight junction degradation, particularly ZO-1 and claudin-5 (Desai et al., 2002; Klein et al., 2019; Lucas et al., 2006). Alteration of the BBB integrity increases the opportunity for the viruses and cytokines to pass the BBB and facilitate the infiltration of periphery immune cells into the CNS, resulting in brain injury and exacerbating neuropsychiatric symptoms (John et al., 2003; Kempuraj et al., 2017).

The E protein may induce the production of inflammatory factors through two potential mechanisms. TLR2 can sense the E protein (Planès et al., 2022; Zheng et al., 2021). The E protein specifically interacts with the TLR2 pathway, activating NF-κB and inducing the release of pro-inflammatory cytokines and inflammatory chemokines. Additionally, it was found that the SARS-CoV envelope protein interacted with syntenin of the host cell. This interaction caused syntenin to be redistributed to the cytoplasm, where it caused the upregulation of inflammatory cytokines (Jimenez-Guardeño et al., 2014). This would cause an exacerbated immune response, leading to tissue damage, edema, and ultimately the characteristic acute respiratory distress syndrome, which was consistent with the histopathological characteristics induced by the SARS-CoV-2 E protein in the mouse spleen and lung (Xia et al., 2021).

TLRs identify pathogen-associated molecular patterns (PAMPs) and trigger immune responses on interaction with infectious pathogens. TLR1–9 are expressed in microglia. The expression of TLRs in microglia is controlled in response to pathogens (McKimmie and Fazakerley, 2005). Microglial activation and neurotoxicity have been connected to TLRs (Jack et al., 2005; Olson and Miller, 2004). Several TLRs (TLR1, TLR2, TLR4, etc.) have an association with disease progression in patients with COVID-19 (Zheng et al., 2021). In particular, TLR2, which can sense the E protein, is necessary for inflammatory responses to coronavirus. We also found that the E protein upregulated TLRs in multiple brain regions and triggered microglial inflammatory response via TLR2 in vitro. TLR2 is crucial for the microglia in response to viruses (Aravalli et al., 2005). TLR2 activation induced microglia to release NO and other cytotoxic substances through multiple ligands (Ebert et al., 2005).

This study has several limitations. We found that the E protein caused neuroinflammation by activating TLR2 and led to depression and dysosmia. Given that TLR2 receptors are expressed in both astrocytes and neurons, we need to further clarify whether the depression-like behaviors and olfactory dysfunction we observed were caused by microglial activation via TLR2. In the CNS, microglia constantly monitor the microenvironment and produce substances that have an impact on nearby astrocytes and neurons (Gomez-Nicola and Perry, 2015). Despite the fact that the interplay between glial cells and neurons is crucial in brain pathophysiology (Zhang et al., 2016a). It is impossible to disregard the immediate impact of SARS-CoV-2 on neurons. SARS-CoV-2 can infect neurons directly (Beckman et al., 2022; Lyoo et al., 2022). It deserves further exploration whether the E protein causes injury to neurons and the relationship between this effect and the neurological complications of SARS-CoV-2.

## Supporting information

Highlights

## Ethic Approval

The experimental procedures were approved by the Peking University Biomedical Ethics Committee experimental animal ethics branch.

## Conflicts of interest

The authors declare that they have no competing interests.

## Funding

This work was supported by National High Level Hospital Clinical Research Funding (Scientific Research Seed Fund of Peking University First Hospital): 2022SF20 (Recipient: Wenliang Su).

## Notes

### Competing Interest Statement

The authors have declared no competing interest.

