## Supplementary material for "SARS-CoV-2 Envelope protein triggers depression and dysosmia via TLR2 mediated neuroinflammation": Highlights

• SARS-CoV-2 Envelope protein evokes obvious neuroinflammation in central nervous system.

• Inhibiting microglia, but not astrocyte, successfully alleviates depression and dysosmia induced by Envelope protein.

• Blocking TLR2 mitigates depression and dysosmia induced by Envelope protein.
